# Complex roles for a conserved two-component system in a honey bee symbiosis

**DOI:** 10.64898/2026.09.25.754351

**Authors:** Joselyn M. Molinar, Audrey J. Parish, Deauna Johnson, Ansel Trinidad, Danny W. Rice, Jonathan Trinidad, Barry Stein, Ian P. Reynolds, Clay Fuqua, Iren L. G. Newton

## Abstract

Elucidating how microbes associate with their hosts and respond to different environments is fundamental to our understanding of symbiosis. Here, we investigate the molecular and genetic mechanisms enabling the honey bee symbiont *Bombella apis* to colonize and persist in harsh honey bee microenvironments, like royal jelly and the larval gut. By using evolved *B. apis* strains, we demonstrate that the broadly conserved two-component system (TCS) ChvGI is integral for the bacterium’s ability to withstand royal jelly and the larval gut. Furthermore, we show that in *B. apis* ChvGI also regulates critical bacterial behaviors such as motility, flagellar production, and biofilm formation. In other *Alphaproteobacteria,* ChvGI is known as a master regulator that mediates the bacterial responses to stressors and is also found to be involved in symbiosis, often through the production of EPS and biofilm formation. Our work underscores the broader significance of ChvGI as an important TCS in *Alphaproteobacteria* that regulates bacterial behavior in response to different types of stressors and uncovers a genetic mechanism that allows an important honey bee symbiont to survive in diverse host environments.

**Importance:** *Bombella apis* is an alphaproteobacterial honey bee symbiont that provides protection against *Aspergillus* species, fungal pathogens of honey bees, and acts as a larval dietary supplement in nutrient poor conditions. This organism is compelling for both its ability to grow in the presence of antimicrobial royal jelly and in its ability to thrive in various hive and insect-associated environments, including the developing larva. Here, we have implicated the two-component system ChvGI as important for the survival of *B. apis* in royal jelly and for regulating bacterial behaviors like biofilm formation, motility, and host association. ChvGI is a broadly conserved TCS in *Alphaproteobacteria* that is widely known to play an important role in mutualistic and pathogenic symbioses, as well as regulating multiple bacterial behaviors. We hypothesize that ChvGI may aid in transition of *B. apis* through the different host environments it occupies.

## Introduction

The European Honey Bee (*Apis mellifera*) is the most important agricultural pollinator on the planet and hosts a characteristic microbial community that has been well studied (1). Pollination efforts from honey bees contribute 15 billion USD annually to the US economy alone (2) although colonies have been in decline for decades (3). The honey bee worker microbiome consists of key bacterial taxa that are repeatedly sampled from bees across geography (4, 5) and seasons (6). This microbial community provides significant benefits to the host, ranging from nutrient breakdown (7), to pathogen protection (8), and host response to pesticide exposure (9). There is also evidence that these bacteria associated with the worker bee might play a role in nest mate recognition and behavior (10–12). Although the worker microbiome is well studied, we know comparatively little about microbes from other castes and hive environments. Honey bees are eusocial and reside within a built environment (the hive) with tens of thousands of sister bees in different developmental stages (the colony). This level of organization means that the colony is the unit of selection, with the queen being the only reproductively capable female. Because of the presence of multiple castes (worker, drone, queen) and developing brood (larvae) as well as foraged resources (pollen and nectar), there are a multitude of environments for microbes to occupy. Therefore, due to the eusocial nature of the insect, it is important to recognize the microbes present in different colony and hive environments and the benefits they might provide to the bee. One critical determinant of these microbial environments is royal jelly, an acidic and viscous glandular secretion produced by nurse bees that contains antimicrobial peptides, creating an environment recalcitrant to microbial growth (13). Therefore, although many different microbes are associated with the honey bee, attributes of different niches in the hive and colony, like the presence of royal jelly, may determine which microbes occupy these environments.

*Bombella apis* is a low abundance bacterial community member in the worker hind gut but is known to occupy other important environments in the colony (the queen gut and larvae) and can survive in the presence of royal jelly. *B. apis* was not previously considered a core microbiome member of the honey bee worker gut (14), but identified in larval association (15) and in honey bee queens (16–19) using 16S amplicon approaches, microbiological plating, as well as metagenomics. Other hive environments in which *B. apis* is found include the worker hypopharyngeal gland, where royal jelly is produced, and the crop. These environments all have in common the presence of royal jelly (**Figure 1**). Therefore, *B. apis* must be able to tolerate the antimicrobial activity of royal jelly. Indeed, using *in vitro* assays and different concentrations of royal jelly, it is evident that various *B. apis* strains can persist while other larval associated bacteria are inhibited (20). *B. apis* in turn provides significant benefits to the honey bee. For example, in nutrient poor conditions, *B. apis* supplements bee larvae nutrition, through the production of lysine, an important essential amino acid (20). Additionally, *B. apis* provides protection against species of *Aspergillus,* known honey bee fungal pathogens, by secreting an active antifungal molecule (21).

**Figure 1.**
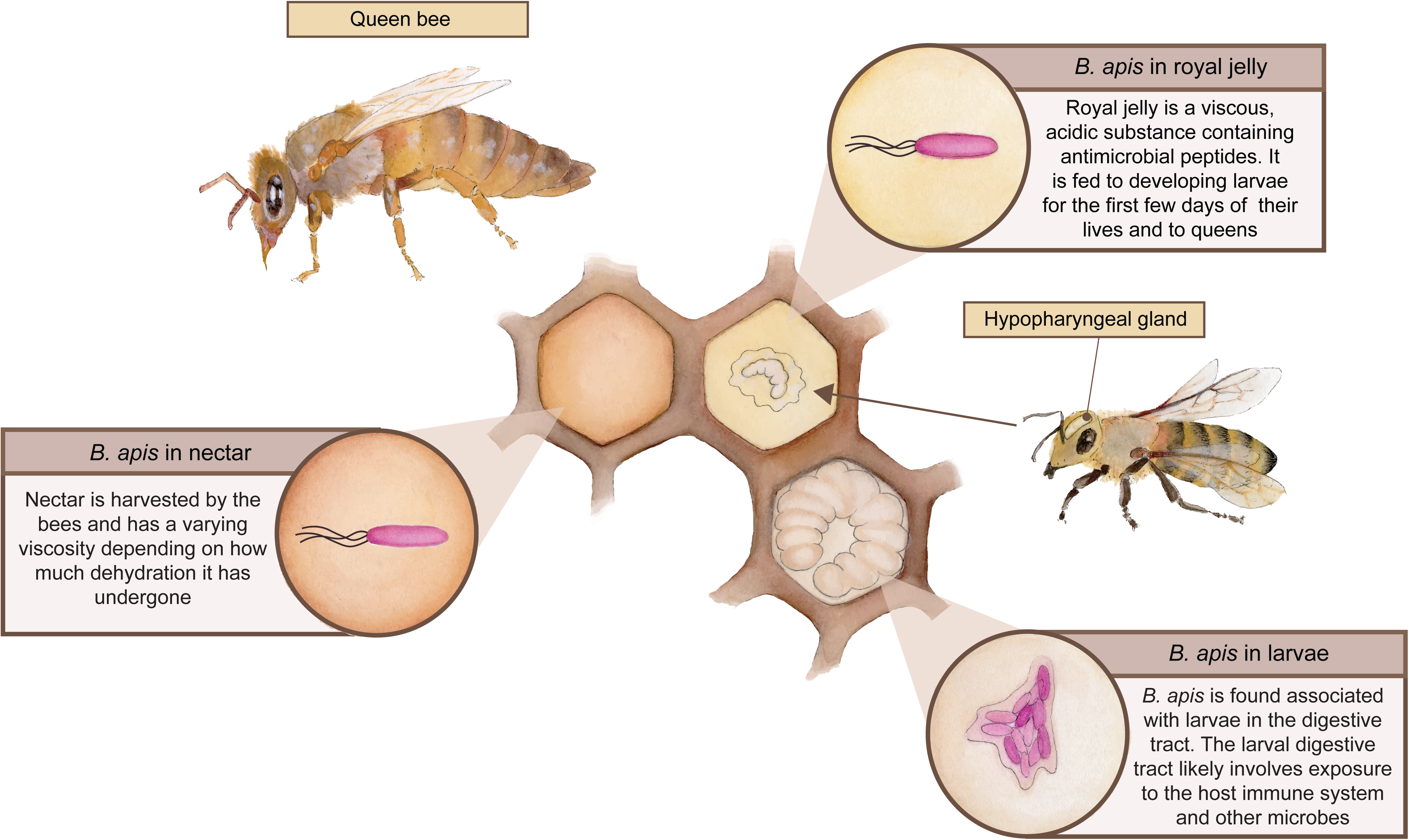
The honey bee microenvironments *Bombella apis* occupies and the behaviors potentially important for its survival. *B. apis* can be found in the nectar and food stores processed in the hive, which can be of varying viscosity depending on how much dehydration they have undergone; we hypothesize that motility may be important for *B. apis* to move through this environment and/or acquire oxygen. *B. apis* can also be found in the queen and larval guts, where we hypothesize that biofilm formation may help *B. apis* avoid exposure to specific host stressors like antimicrobial peptides or the host immune system. Finally, *B. apis* may also be found in royal jelly, a semifluid and acidic substance containing antimicrobial peptides that is produced in the hypopharyngeal glands of nurse bees. Royal jelly is fed to larvae for the first few days of their lives and to the queen. Here, we hypothesize that motility may also be important to survive this incredibly viscous environment. (Artwork by Aya McGee)

Honey bees use trophallaxis to exchange gut contents after foraging to feed the queen and developing larvae, to make honey from foraged resources and bee bread, the product of fermented bee pollen and honey (22). This regurgitation – mouth to mouth – allows for microbial exchange as well and is likely one way in which *B. apis* is passed between different colony environments. These processes suggest that several factors determine the distribution and density of *B. apis* across the colony landscape, including the presence of royal jelly and whether *B. apis* was trafficked through these different niches through trophallaxis.

Bacteria are frequently exposed to stressful conditions during one or more stages of host-association (23). One of the best studied animal symbioses is that of *Vibrio fischeri* and *Euprymna scolopes* (Hawaiian Bobtail Squid), in which the free-living bacteria colonize the light organ of the squid host. Establishment and maintenance of the *V. fisheri* monoculture in the light organ is a highly selective process, referred to as the “winnowing” (24). Colonizing *V. fischeri* are subjected to acidic pH, reactive oxygen species (ROS), nitrous oxide, and antimicrobial peptides (AMPs) and in turn mount a substantial stress resistance response including exopolysaccharide (EPS) production, outer membrane modification, and pH adaptation (25–27). For the nematode association of *Xenorhabdus* species the bacteria face stressful competitive interactions via production of bacteriocins (28). Additionally, a *Xenorhabdus* CRISPR system is required for symbiosis, suggesting the presence of bacteriophage (29). In plant-associated bacteria, host-induced stresses such as toxic exudates, low pH, ROS, and AMPs including defense responses, play an important role (23). Association of symbiotic rhizobia with plants and recognition of outer membrane LPS causes an oxidative burst by legumes, that is tolerated in part by exopolysaccharide production to initiate formation of nitrogen-fixing nodules (30). Nodulation by *Sinorhizobium meliloti* and other rhizobia is in part shaped by a subclass of host-produced, cysteine-rich plant defense AMPs (31), which cause outer membrane stress in a wide range of bacteria. Acidic pH at the sites of plant infection is also a common stress to which symbiotic bacteria must acclimate. In *S. meliloti* an related Alphaproteobacteria, like the plant symbiont *Agrobacterium tumefaciens*, the ChvG-ChvI two component system (TCS) functions to activate responses to low pH and outer membrane stress (32), often controlling the expression of a wide array of genes including outer membrane proteins, exopolysaccharides, and motility (33–35).

Here, we examined the molecular mechanisms that contribute to *B. apis’* ability to survive the stressful conditions of host association using experimental evolution (EE) through laboratory passage in royal jelly. We identified consistent mutations across the populations recovered from royal jelly in a region encoding homologues to the ChvG-ChvI TCS. We then isolated several ChvG mutant lineages from these populations and performed phenotypic assays to determine the impact of these mutations (**Figure 2**). ChvG mutants were found to have an increase in swimming motility and royal jelly persistence with decreased ability for biofilm formation and larval host association. Since ChvG is a TCS sensor kinase that acts with ChvI to regulate transcription of gene networks in multiple Alphaproteobacteria, we used RNA-seq to measure differential expression resulting from mutations in ChvG, and our results suggest changes in outer membrane formation and stability as well as purine ribonucleoside triphosphate metabolic processes. Finally, electron microscopy (EM) and mass spectrometry revealed differences in flagella produced when comparing wild type and a ChvG mutant *B. apis* strain. Overall, our work identifies ChvG as an important molecular determinant for *B. apis* acclimation to honey bee microenvironments.

**Figure 2.**
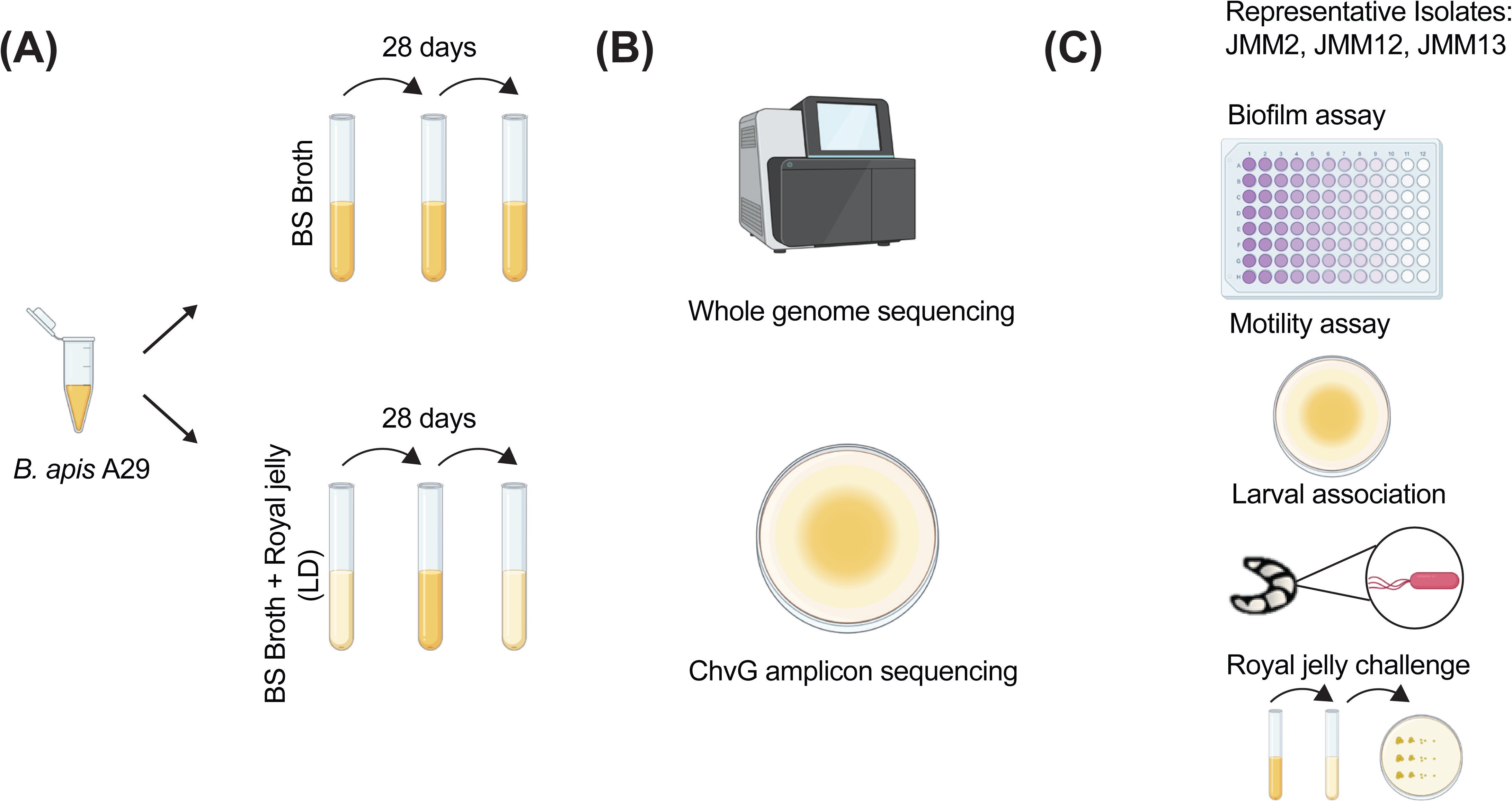
Schematic outline of experimental design for in lab evolution of *B. apis* strain A29 and phenotypic typing. (A) To identify the mechanisms in which *B. apis* can survive in royal jelly we first experimentally evolved *B. apis* A29 in parallel in either BS broth as a control and comparator and in BS broth supplemented with 50% royal jelly (similar to larval diet used to rear larvae in laboratory conditions). Each condition contained 10 independent populations to briefly assess parallel evolution, and the populations evolved in larval diet were alternated with passages in just BS broth to ensure the recovery of CFUs. (B) After 28 days of daily passaging, all populations were sent for sequencing and a subsequent Breseq analysis was used to identify any acquired mutations. Populations were then streaked for isolation and the ChvG region was amplified and sent for Sanger sequencing to confirm mutations. (C) Isolated mutants JMM2, JMM12, and JMM13 were then tested for specific phenotypic assays that align with typical behaviors ChvGI is known to control, as well as verifying that these mutations allow for enhanced survival in bee-specific environments.

## Results

### Experimental evolution of *B. apis* A29 in royal jelly reveals mutations in loci homologous to ChvGI

To identify genetic determinants in *B. apis* that contribute to growth in royal jelly, we performed EE on progenitor strain A29. Currently, we are unable to introduce DNA into *B. apis*, leading us to leverage the power of forward genetics. A confluent culture of *B. apis* was divided into two conditions, each with 10 replicates to assess the effects of parallel evolution. The control condition consisted of passage through standard liquid media used to grow *B. apis* (BS broth) while the experimental condition alternated plain BS broth and BS broth supplemented with royal jelly to a 50% concentration intended to mimic the larval diet (LD) (see materials and methods for more detail). After 28 daily passages, entire populations were sequenced, as well as the progenitor strain, to determine which mutations had been selected for in each condition (**Table S1**). The *B. apis* A29 genome was sequenced using Oxford Nanopore Technology and published short reads were used to polish the sequences resulting in a single, closed chromosome of 2.02 Mbp. All passaged populations were sequenced using Illumina short reads, which were mapped to the complete *B. apis* A29 assembly using Breseq to identify mutations (see *Data Availability* statement for all accessions). Every population passaged through royal jelly, and none of the control populations, developed mutations in a region of the genome containing homologs for ChvGI, a histidine kinase – response regulator pair conserved in alphaproteobacteria (**Figure 3A, B**). Mutations in the histidine kinase *chvG* were uniformly insertions or deletions predicted to result in loss of function for the protein, whereas mutations in *chvI* included multiple point mutations. The *B. apis* ChvG periplasmic domain diverges from those ChvG homologs that interact with ExoR, but its architecture and cytoplasmic domain structure is quite consistent with this group of proteins. The cytoplasmic portion of the protein has the canonical HAMP and HisKA domains, as well as a more C-terminal ATPase domain (**Figure S1A**). ChvG is thought to phosphorylate the response regulator ChvI, and phosphor-ChvI is activated for DNA binding. ChvI from *B. apis* is well conserved with other ChvI orthologs and has the canonical two domain structure, with an N-terminal phosphor-accepting receiver domain and a C-terminal DNA binding domain (**Figure S1B**). All of the *B. apis chvI* mutations identified after royal jelly passaging occurred in either the hinge or C-terminal DNA binding domain. Interestingly, four of the six of these mutated an arginine residue to an uncharged residue (**Figure 3A**).

**Figure 3.**
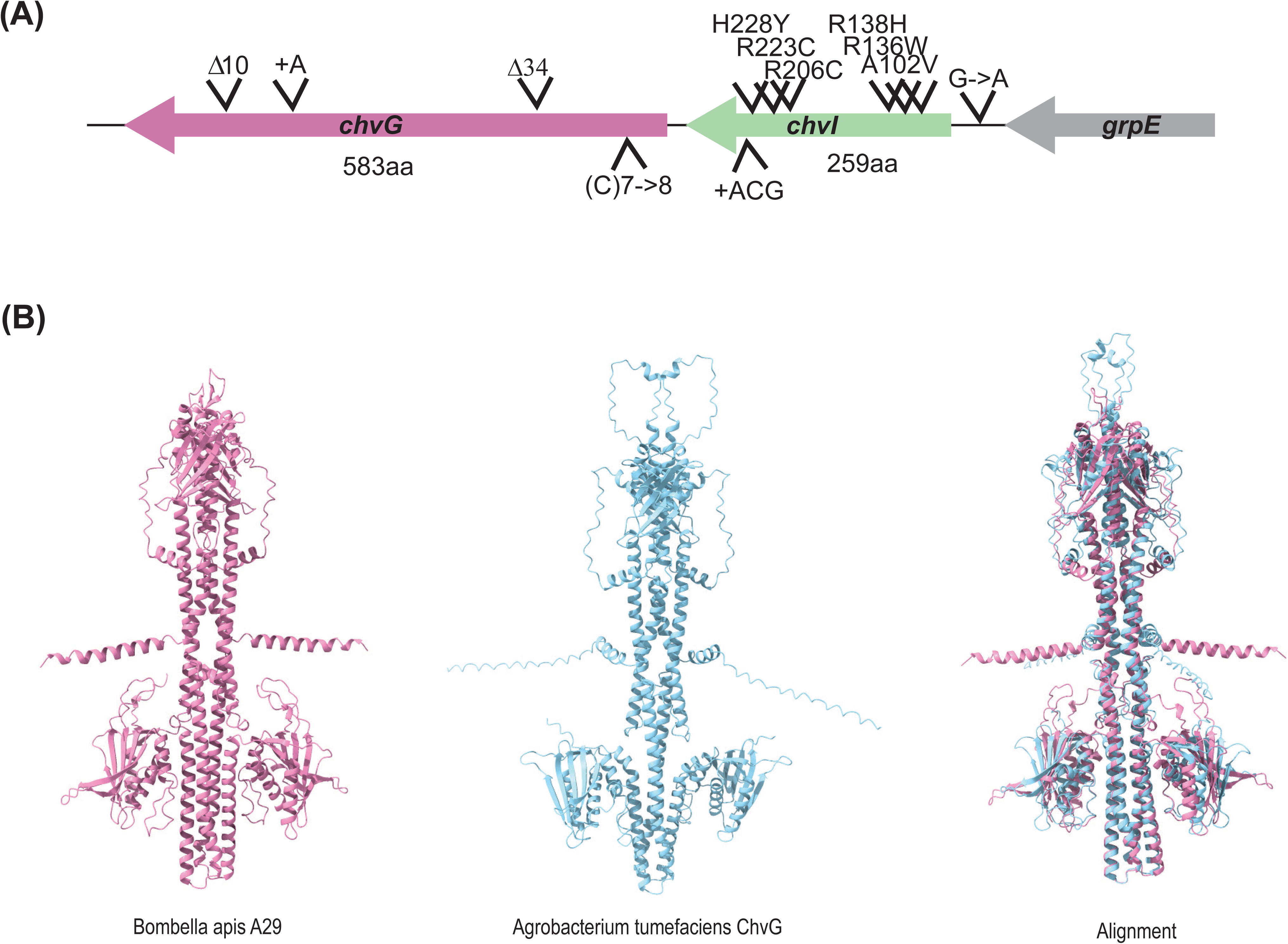
*B. apis* ChvGI sensor histidine kinase and response regulator pair are selected for under royal jelly evolution. (A) Schematic representations of mutations found across resequenced *B. apis* populations passaged in royal jelly. Mutations in *chvG* abrogate protein production. (B) Comparison of *B. apis* and *Agrobacterium tumefaciens* ChvG structures generated using Alphafold (Abramson et al., 2024) and ChimeraX (Meng et al., 2023). The RMSD across 162 pruned residues was 1.24Å.

### ChvG mutation impacts phenotypes important for host environments

To determine the impact of ChvGI on *B. apis* behavior, we isolated ChvG null mutants (JMM2, JMM12, and JMM13) from the EE LD populations and confirmed that they did harbor mutations in ChvG using PCR amplification of the gene and chain-termination sequencing. Since evolution in royal jelly consistently resulted in mutations within the *chvGI* region we first assessed the ability of the individual isolated *chvG* mutants to tolerate and survive in royal jelly compared to the ancestral strain A29. ChvG mutants and A29 were normalized to mid exponential phase (OD_600_ = 0.4) and back-diluted into 30% royal jelly, with CFUs checked in triplicate at 6-hour intervals over the course of 24 hours. The most prominent observation is that the A29 strain does not grow in royal jelly and instead we see a loss of CFUs consistently over the time course (growth rate= –0.04). In contrast, after 6 hours, all 3 *chvG* mutants increase in abundance with an average growth rate of 0.51. Additionally, all mutant strains remain at high viable counts until 12 hours, when they start to decline, and reach an average maximum yield that is higher than the maximum yield reached by progenitor A29 (3.1×10^8^ compared to 1.3×10^7^, respectively) representing an average fold difference of 23.3 (Range: 16.25-32.5). These growth dynamics suggest that *chvG* null mutations enhance the ability of *B. apis* to multiply and survive in royal jelly (**Figure 4A**).

**Figure 4.**
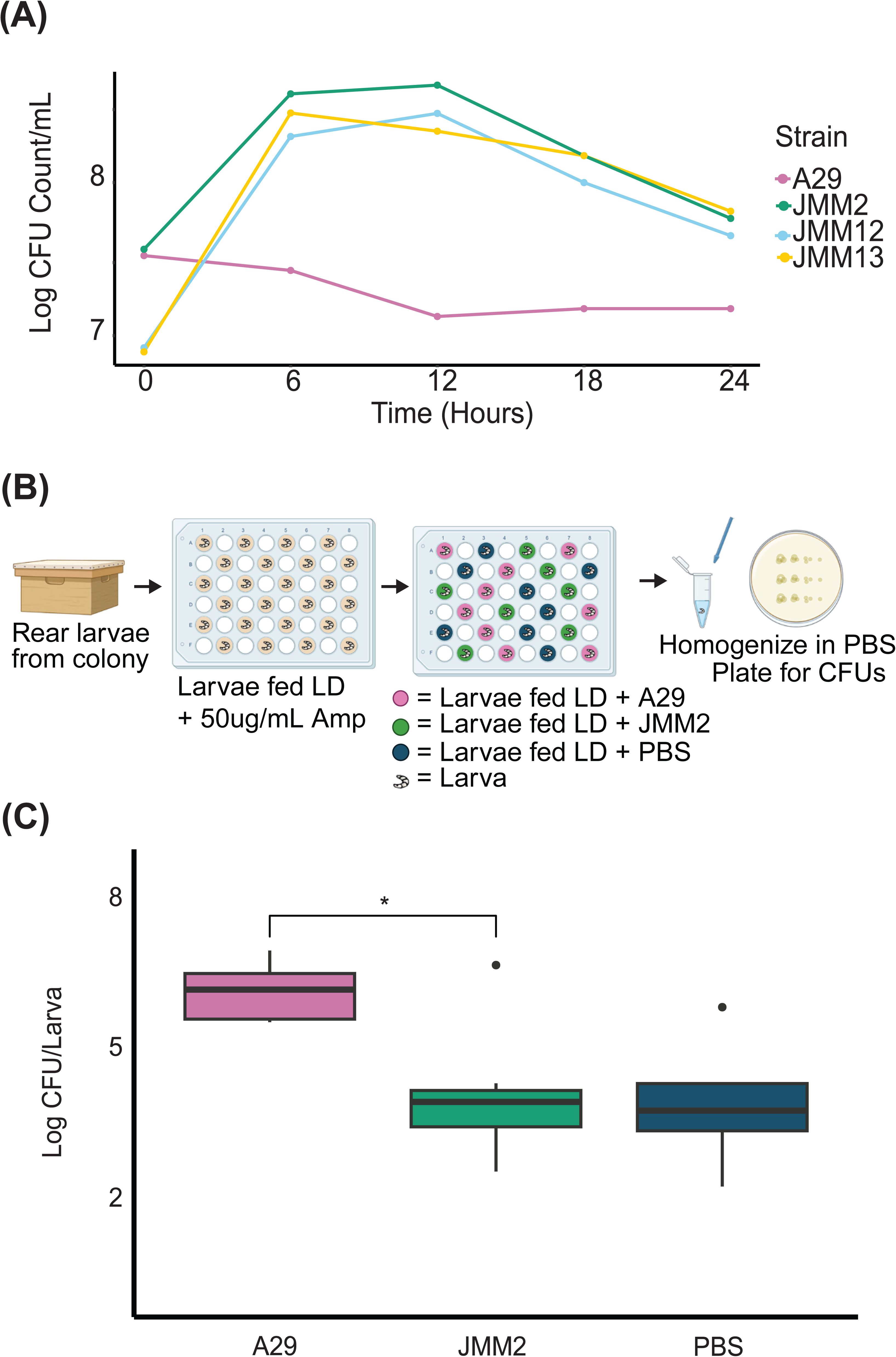
ChvG plays an important role in royal jelly survival and larval colonization in *B. apis.* (A) A 24-hour growth curve in 30% royal jelly uncovers the *chvG* mutants enhanced ability to grow and survive in royal jelly (JMM2: µ=0.39, JMM12: µ=0.53 and JMM13: µ=0.60) compared to progenitor strain A29 (µ=–0.04). (B) Schematic representation of the larval rearing process. Larvae are reared from the colony, placed in queen cups in a 48-well plate and fed larval diet (LD). 24 hours after initial rearing larvae are then fed LD with 50 ug/mL of ampicillin. 48 hours after rearing larvae are fed LD + either *B. apis* strain A29 or JMM2. From then on larvae are fed LD with no additional components. After 6 days, larvae are homogenized in PBS and plated for CFU counts. (C) When looking at larval association compared to A29, JMM2 has a significantly decreased ability to colonize larvae (Kruskal-Wallis; *χ²* = 8.7, df = 2, *p* = 0.01266; Wilcoxon rank-sum pairwise comparison of JMM2 to A29 with Benjamini-Hochberg correction: A29 (*n*=6) and JMM2 (*n*=8), *W* =44, *padj* = 0.035)

Considering the locations *B. apis* can be found in the hive (**Figure 1**), coupled with the known impact ChvGI has on host association in other organisms, we were curious whether the *B. apis chvG* mutant would be impacted for its ability to colonize larvae relative to the parent strain A29. Therefore, we tested the ability of the *chvG* mutant JMM2 and A29 to colonize larvae through an *in vitro* rearing assay. Briefly, first instar larvae (larvae that are early in their developmental stage) were grafted from hives located in our local apiary, placed in sterile multiwell plates containing queen cups (small plastic vessels used for queen rearing) and fed UV sterilized *in vitro* larval rearing diet (LD) adapted from Parish et al. 2022 (see methods). After 24 hours, larvae were fed LD supplemented with 50 ug/mL of ampicillin to reduce the abundance of microbes that the larvae had acquired in the hives. At the 48-hour mark, larvae were fed LD supplemented with a solution of 10^5^ CFU/mL of either A29 or JMM2, or an equal volume of sterile PBS as a control. Larvae were then fed LD daily in accordance with the feeding schedule outlined in the methods. At the end of 6 days and before pupation, larvae were surface sterilized to remove surface microbes, homogenized in PBS, and subjected to viable counting (CFUs) on BS plates (**Figure 4B**). JMM2 shows a decreased ability to colonize larvae compared to A29 (Kruskal-Wallis: χ^2^ = 8.7, df = 2, *p* = 0.01266; Wilcoxon rank-sum pairwise comparison of JMM2 to control A29 with a Benjamini-Hochberg correction: A29 (*n*=6) and JMM2 (*n*=8), *W =* 44, *padj*= 0.035). The amount of microbial growth recovered from the JMM2 inoculation is not statistically different from the PBS control suggesting that this growth is likely representative of the bacteria acquired from the natural hive environment prior to the laboratory incubation of the larvae (**Figure 4C**). Overall, these results indicate that a *chvG* null mutant has a decreased ability to colonize larvae compared to the progenitor strain, highlighting the importance of a functional ChvG for host colonization.

### *B. apis* strains with ChvG mutations show decreased biofilm formation

In other *Alphaproteobacteria,* such as *A. tumefaciens,* ChvGI is known to regulate biofilm formation (32). Additionally, biofilm formation is typically important as bacteria associate with the host and aid in withstanding host-associated stressors (36). Therefore, we tested the biofilm formation ability of three of the different isolated *chvG* null mutants compared to the progenitor strain, A29, in a static 96-well plate biofilm assay. Biofilm formation in all strains was assessed by quantifying the biomass after 24 hours using crystal violet and normalizing culture density (OD_600_) to account for variable growth. *B. apis* strain A29 forms robust biofilms and boasts a score that is nearly two times higher than all three of the *chvG* mutants that show undetectable biofilm formation. Overall, we observed a statistically significant difference amongst strains with regards to biofilm formation (F=116.5; df = 3; *P* = <0.001) (**Figure 5**), with a Tukey’s HSD pairwise comparison to A29 revealing that biofilm formation of each ChvG mutant was significantly decreased (JMM2 (*Mdiff* = –1.52, 95% CI [-1.79, –1.24], *p* < 0.0001); JMM12 (*Mdiff* = –1.49, 95% CI [-1.77, –1.21], *p* < 0.0001); JMM13 (*Mdiff* = –1.63, 95% CI [-1.90, –1.35], *p* < 0.0001), demonstrating that mutation of *chvG* results in a nearly 2-fold significant decrease in biofilm formation. Therefore, we conclude that ChvG plays an important role in biofilm formation in *B. apis*.

**Figure 5.**
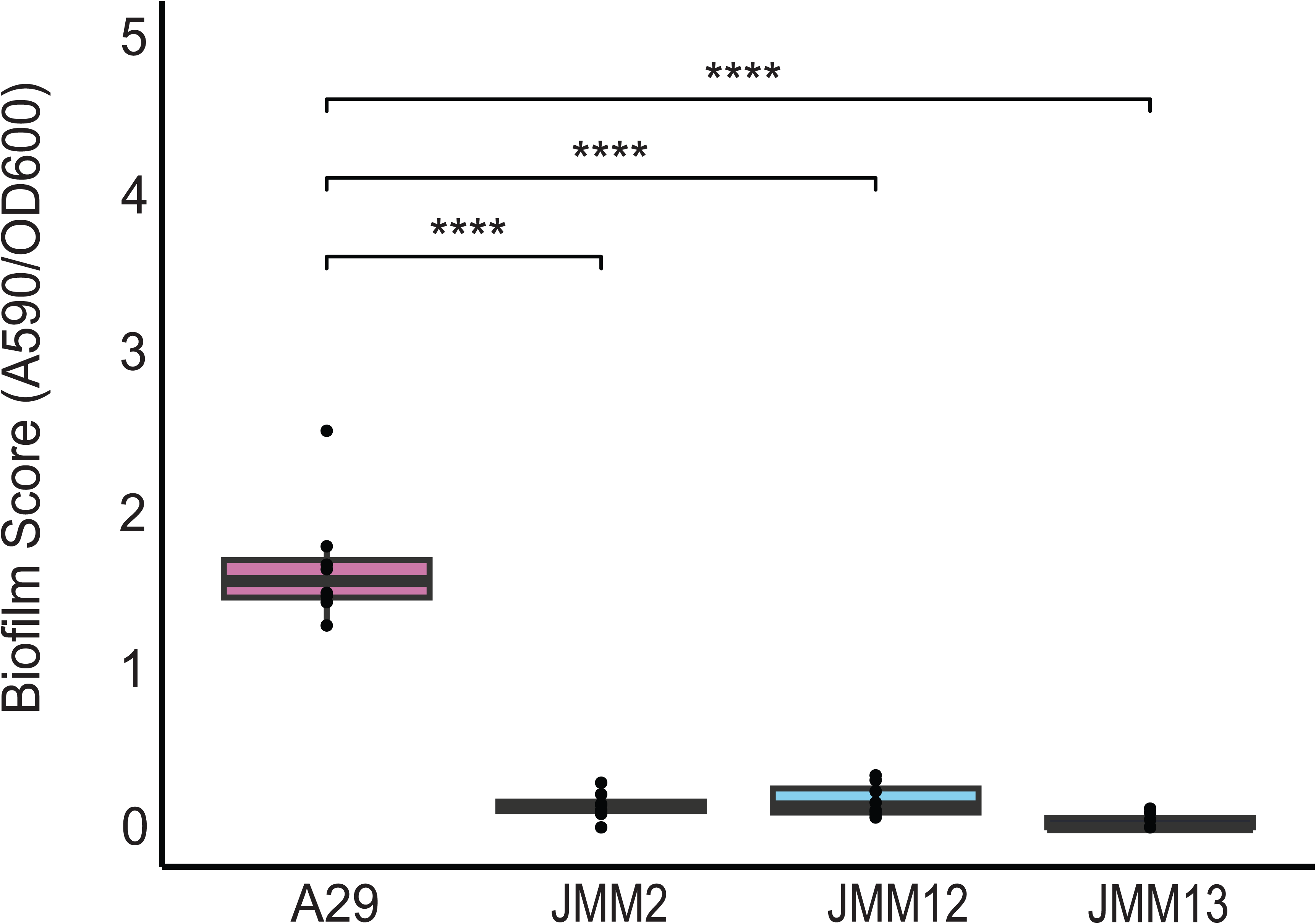
ChvG is integral for biofilm formation in *B. apis* Biofilm assay reveals a clear decrease in biofilm formation ability for all ChvG mutants compared to progenitor strain A29. (One-way ANOVA; F(3,28)=116.5, *P* = <0.001; Tukey’s HSD pairwise comparison to A29 revealed that JMM2 had significantly decreased ability to form biofilms (*Mdiff* = –1.52, 95% Cl [-1.79, –1.24], *p* < 0.0001); JMM12 (*Mdiff* = –1.49, 95% Cl [-1.77, –1.21], *p* < 0.0001); JMM13 (*Mdiff* = –1.63, 95% Cl [-1.90, –1.35], *p* < 0.0001.

### A ChvG mutation leads to increased swimming behavior and heightened production of flagellar filaments

It is well established that motility heavily impacts a bacterium’s ability to navigate towards or away from specific stimuli, serving as a critical behavior in model symbioses. Furthermore, the ChvGI system in other *Alphaproteobacteria* is responsible for regulating motility. Therefore, we hypothesized that a *B. apis* ChvG null mutant might also impact swimming behavior. We next qualitatively assessed the swimming ability of the *chvG* null mutants compared to A29. Each strain was inoculated into the center of a 0.2% BS agar plate and assessed after 24 hours. The ChvG null mutants show enhanced motility compared to A29, both in terms of swim radius and speed, suggesting a genetic link between ChvG and motility in *B. apis* (**Figure 6A**). As shown in figure 6A, the progenitor strain A29 shows growth in the center at the point of inoculation but does not display any outward growth that would indicate swimming behavior. If left longer than 24 hours, swimming motility will begin to show (**Figure S2**). Additionally, depending on inoculation, A29 will display sliding motility where it passively moves on top of the agar surface through the production of a surfactant and dendrites are produced (37) (**Figure S2**). In contrast, all 3 ChvG mutants show enhanced swimming motility at 24 hours, where we see JMM12 and JMM13 produce swim rings that nearly encompass the entire diameter of the petri dish. JMM2 also shows enhanced motility and a swim radius that is much larger than the progenitor strain but does not extend as far as the other two mutants. This is likely a result of inoculation differences as this assay has been repeated multiple times and JMM2 has also formed swim rings that occupy the entire petri dish at the same rate as the other mutants. Observing these striking differences in motility between the *chvG* mutants and the progenitor strain A29, we were then motivated to evaluate swimming behavior at the single cell level using microscopy. Cultures of all strains were grown to mid-exponential phase and observed at 100X magnification. There is a clear difference between progenitor A29 and ChvG mutant JMM2. In the A29 sample the cells are predominantly seen in aggregates or twitching in place, with a small fraction of swimming cells. In stark contrast, most of the cells visible in the JMM2 movie can be seen swimming or tethering in place, potentially where the flagellum attached to the surface (**Supplemental Movies**). The striking difference in the number of cells swimming in these samples further supports our hypothesis that ChvGI plays an important role in regulating motility in *B. apis*.

**Figure 6.**
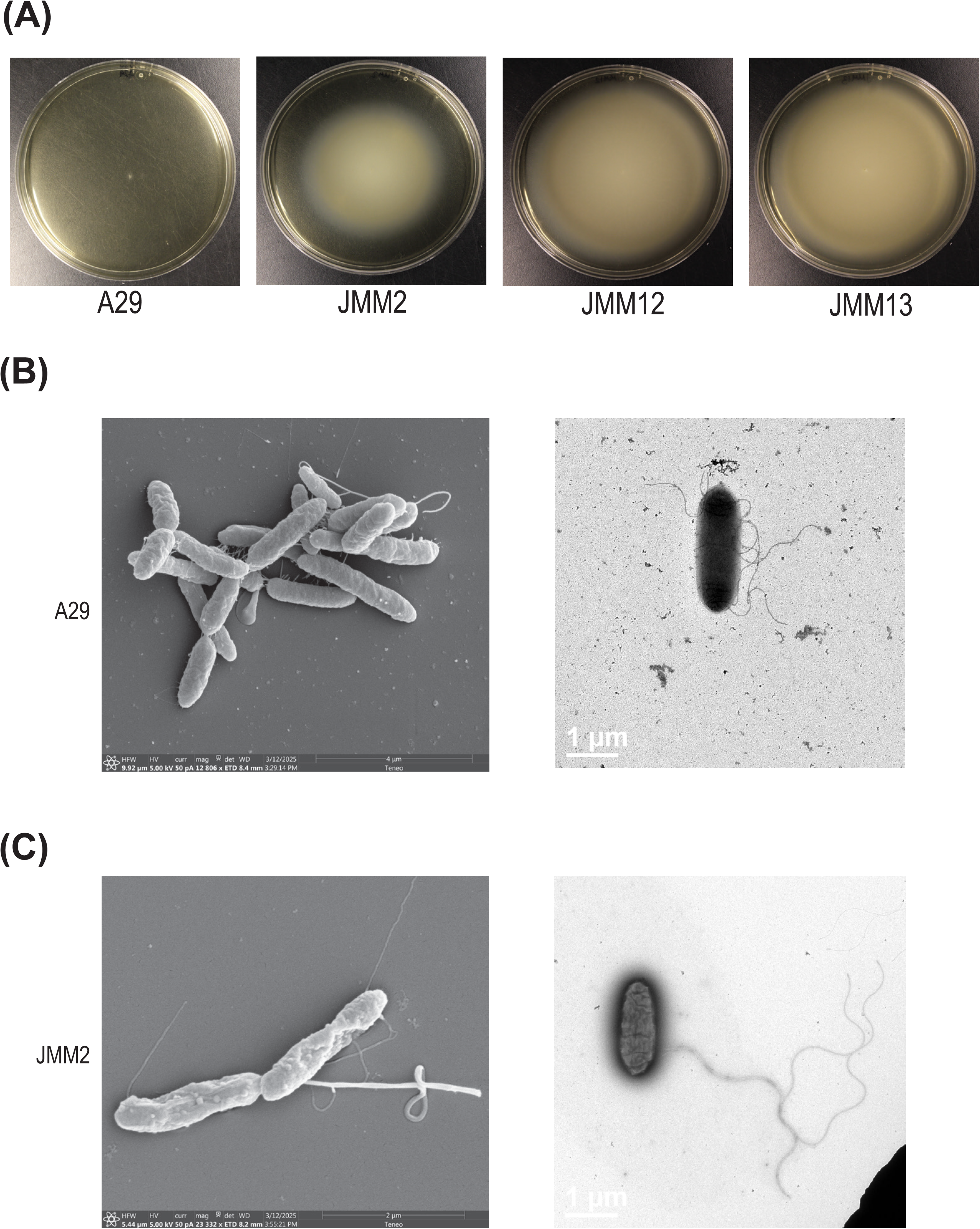
ChvG regulates motility and impacts flagellar production. (A) Swim behavior assessed by inoculating each respective strain into the center of a 0.2% agar plate revealed that all 3 *chvG* mutants show enhanced motility, regarding the swim radius and speed, compared to progenitor A29. (B) Scanning electron microscopy (SEM) image (left) shows A29 in aggregates with some bacteria producing flagella. A representative transmission electron microscopy (TEM) image (right) shows a single A29 cell with thinner flagellar filaments. (C) A SEM image (left) of ChvG mutant JMM2 shows a thick protruding appendage. A TEM image (right) of JMM2 shows a single cell with a thicker flagellar filament compared to A29.

The enhanced swimming behavior displayed suggests that motility may be an important phenotype in the royal jelly environment and that a *chvG* null mutation might alter either the type, number, or form of the flagella used by *B. apis*. We therefore took three approaches to quantitatively assess differences in flagella between A29 and one *chvG* mutant, JMM2. First, we used scanning electron microscopy (SEM) and transmission electron microscopy (TEM) to visualize the flagellar filaments. Micrographs of both strains reveal flagellar filaments but also, we observed thick protruding structures in JMM2 in addition to a much longer flagellar filament (**Figure 6C**). Interestingly, these structures are much thicker than typical flagella and not observed in the A29 images (**Figure 6B**). We wondered what these structures might be and if they were made of flagellin. The A29 genome contains 2 flagellin homologs and we therefore reasoned that differences in flagellar structure could be due to different monomers in the filament. To address this question, we turned to mass spectrometry.

Mass spectrometry can be used to quantify the relative abundance of different flagellins (38). Briefly, we individually cultured both JMM2 and A29 to mid exponential phase when flagella production should be heightened, centrifuged the bacteria so flagellar filaments would be sheered from the cell body and remain in the supernatant, then pelleted the flagella via ultracentrifugation. The flagellar preparations were digested with trypsin and analyzed via liquid chromatography-mass spectrometry. In the individual runs, we identified peptides matching both flagellin homologs in JMM2, but only 1 flagellin was identified in A29 (**Table 1, Table S2**) although the abundance of both was higher in JMM2 (with a ratio of 1.9 for area under the curve when comparing JMM2/A29 for the flagellin identified in both individual runs). Strikingly, in progenitor strain A29, there are no other flagellar-related proteins identified in the individual run, suggesting that they are either not present in a high enough abundance to be confidently identified, or are broken off during the flagellar preparation. Conversely, the *chvG* mutant JMM2 has many flagellar-related proteins present in the individual run, namely FlgK, FlgG and FlgN.

**Table 1.** Mass spectrometry results of Bombella apis A29 and mutant JMM2 showing percent coverage, number of total unique peptides and area under the curve.

| Gene description | A29 |  |  | JMM2 |  |  | Accession |
| --- | --- | --- | --- | --- | --- | --- | --- |
|  | % coverage | Num. unique peptides | AUC | % coverage | Num. unique peptides | AUC |  |
| Flagellar basal body rod protein FlgC | - | - | - | 7 | 1 | 4.89E+06 | BIDLMP_00505 |
| Flagellar basal-body MS-ring/collar protein FliF | - | - | - | 6 | 2 | 1.60E+07 | BIDLMP_00543 |
| Flagellar basal-body rod protein FlgG | - | - | - | 19 | 3 | 1.78E+07 | BIDLMP_00549 |
| Flagellar hook protein FlgE | - | - | - | 7 | 2 | 3.12E+07 | BIDLMP_00513 |
| Flagellar hook-associated protein FlgK | - | - | - | 33 | 8 | 4.17E+07 | BIDLMP_00551 |
| Flagellar hook-basal body complex protein FliE | - | - | - | 8 | 1 | LOD | BIDLMP_00509 |
| Flagellar motor stator protein MotA | - | - | - | 8 | 1 | LOD | BIDLMP_00521 |
| Flagellar motor switch protein FliG | - | - | - | 3 | 1 | 3.90E+06 | BIDLMP_00542 |
| Flagellar protein export ATPase FliI | - | - | - | 2 | 1 | 4.70E+06 | BIDLMP_00525 |
| Flagellar protein FlbB | - | - | - | 4 | 1 | LOD | BIDLMP_00504 |
| Flagellar protein FlgN | - | - | - | 7 | 1 | 6.95E+06 | BIDLMP_00529 |
| Flagellar protein FliL | - | - | - | 26 | 2 | 1.87E+07 | BIDLMP_00561 |
| Flagellin | 16 | 2 | 7.58E+08 | 22 | 4 | 1.46E+09 | BIDLMP_00558 |
| Flagellin | - | - | - | 19 | 2 | 1.01E+08 | BIDLMP_00188 |
| Flagellin assembly protein | - | - | - | 20 | 2 | LOD | BIDLMP_00518 |

### ChvG Mutation Significantly Impacts a Specific Gene Network in the B. apis Transcriptome

Given that ChvG functions as the histidine kinase in the canonical TCS ChvGI that regulates transcription in other bacteria, we were curious as to how strain JMM2, with a nonfunctional ChvG and noticeably different phenotypes, would impact the transcriptome compared to progenitor strain A29. Both strains were grown to mid exponential phase in BS media and RNA was harvested for sequencing. Resulting reads were quality controlled and mapped to the complete A29 genome. Using an adjusted P-value of <0.05 we identified 96 genes whose transcription was significantly altered in a ChvG null background.

These 96 genes were further characterized by identifying Clusters of Orthologous Groups (COGs), with roughly half (54) of genes falling into specific categories of different recognized functions (**Figure 7A**) but multiple (42 genes) of unknown function. For those genes that matched an annotated function, COG analysis revealed an overall trend of downregulation in the mutant strain, notably in genes related to coenzyme transport and metabolism, cell wall/membrane/envelope biogenesis, and posttranslational modification/protein turnover/chaperones. This same COG group analysis also identified an upregulation, in the mutant strain, of genes related to energy production and conversion, amino acid transport and metabolism, and defense mechanisms. Considering this mutant was selected for after the continuous passage through royal jelly, we reason that *B. apis* may respond to royal jelly by altering the cell wall and changing nutrient uptake and typical metabolism function.

**Figure 7.**
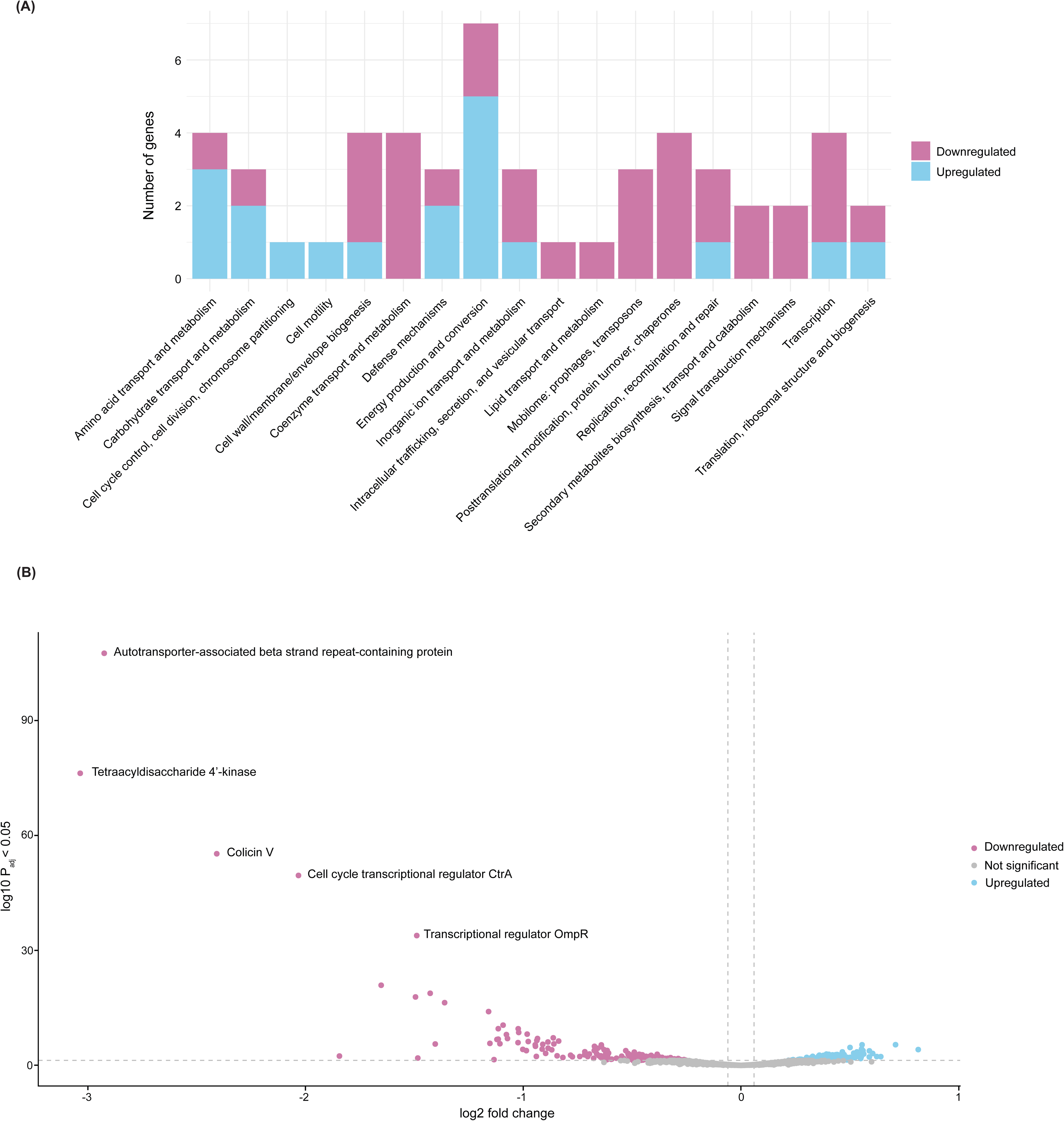
RNAseq of ChvG mutant reveals a general trend of downregulation and significant impact on 96 loci. (A) Clusters of Orthologous Groups (COG) analysis was able to determine a classification for 54 genes with roughly half (42) of genes falling into categories of unknown function (not shown here; see Supplementary Table 2 for full RNAseq results). There is an overall trend of downregulation (shown in pink), with fewer groups showing a pattern of upregulation (shown in blue). (B) Volcano plot displaying the entire transcriptome of JMM2 compared to A29. Insignificant gene comparisons are shown in gray, with significantly downregulated genes shown in pink and significantly upregulated genes in blue. The top 5 significantly downregulated genes are labeled.

In order of significance (**Figure 7B**), the most downregulated gene in the *chvG* mutant, with a 7.5-fold decrease, is annotated as an autotransporter-associated beta strand repeat-containing protein, typically associated with virulence factors (39). The second most significantly downregulated gene with an 8-fold down regulation is predicted to be tetraacyldisaccharide 4’ kinase, a gene involved in the synthesis of lipid A, a component of the lipopolysaccharide layer of the outer leaflet. Another virulence factor related gene, colicin V, known to be important for colonization in *E. coli* (40) is the next most significantly downregulated, with a 5.3 fold depletion. Notably, the fourth most downregulated gene is annotated as CtrA (4 –fold decrease), a known master regulator in Alphaproteobacteria responsible for the regulation of cell division, DNA replication, cell differentiation, and other bacterial behaviors including motility (41–43). Interestingly, depletion of CtrA has been shown to cause defects in cell division and lead to elongated cells (44), a phenotype we do not observe here despite the significant downregulation of *ctrA*. Finally, the fifth most downregulated gene is predicted to be an OmpR-type regulator with a fold depletion of 2.8. OmpR-type regulators have been characterized as global regulators that control bacterial behaviors for acid and osmotic stress response, the expression of virulence factors, motility/flagellar expression, and biofilm formation.

However, determining the function of OmpR-type regulators is difficult given their low DNA-binding specificity and the sheer breadth of their regulons (45). Overall, the ChvG mutant illustrates a widespread transcriptomic down regulation compared to the progenitor strain, A29. Exposing both strains to defined environmental stressors, like osmotic stress or specific royal jelly proteins, may evoke a more selective transcriptomic response.

## Discussion

Elucidating how microbes associate with their hosts and respond to different environments is fundamentally important to our understanding of symbiosis. In this work, we evolved *B. apis* strains by passaging through royal jelly, and identified mutations in *chvG* and *chvI*, that implicated the role of the ChvGI TCS in regulating *B. apis* interactions with the honey bee. We hypothesize that ChvGI acts to acclimate *B. apis* to specific bee environments including royal jelly and the larval gut and regulates specific bacterial behaviors such as biofilm formation and motility. We also evaluated the transcriptomic impact of a *chvG* null mutant, particularly the downregulation of genes related to coenzyme transport and metabolism, cell wall/membrane/envelope biogenesis, and posttranslational modification/protein turnover/chaperones, along with the upregulation of genes related to energy production and conversion, amino acid transport and metabolism, and defense mechanisms.

Still, the complex eusocial nature of honey bees coupled with the intricate built environment of the hive leaves us with many lingering questions. Currently, there is no clear framework describing how *B. apis* is transmitted through the multiple hive environments it has been found in (**Figure 1**), and we hypothesize that the microbial exchange that happens during trophallaxis may be a key factor. In this simulated royal jelly environment, our conditions suggest a fitness advantage for a null *chvG*, given that this was selected for across 10 independently evolved populations. This null mutation derepresses motility, however, we hypothesize that there are conditions and specific environments where motility is regulated as a result of ChvGI activity, potentially through downregulation of the phosphotransfer through the pathway. Given this fitness advantage conferred by a null *chvG* in our royal jelly selection, we predict that ChvGI is very important in other hive environments. If this TCS in a functional state led to general fitness disadvantages across the honey bee association, mutants would have been selected for in other environments, such as when associating with larvae. Despite analyzing many genomes of *B. apis* isolates from honey bee hives, null mutations in *chvGI* have only been observed after our continuous selective regime in royal jelly. Indeed, the larval rearing experiment performed here indicates that a functional *chvG* is integral for *B. apis* to associate with larvae, underscoring its importance in other environments. Additionally, in a subset of populations evolved in royal jelly we noted fixed populations containing insertions in a polymeric stretch in *chvG* (C_7→8_) **(Table S1,** populations 4 and 6). Throughout the process of isolating mutants from these populations, we observed phenotypic indication of a null *chvG*, yet the mutants reverted quickly. This leads us to consider whether ChvGI may function as a contingency locus, allowing *B. apis* to adjust to the different environments it occupies (46). If so, this could explain the varying fitness advantages conferred by ChvGI depending on the specific environment, and would be an example of antagonistic pleiotropy, where a gene may be beneficial or harmful depending on the context (47).

### Significance of experimental evolution in royal jelly

EE is a powerful approach to identifying loci of import in a controlled environment (48). Here, we leveraged EE to probe the mechanisms *B. apis* could withstand and persist in the antimicrobial properties of royal jelly. Across ten independently evolved populations, mutations arose in the *chvGI* operon. The mutations observed in this gene cluster would first have to survive a population bottleneck (due to the serial dilution) and resultant genetic drift (random changes in allele frequencies over time), and selection. As observed in other EE experiments, many naturally occurring mutations are often lost due to population bottlenecks and genetic drift during the daily transfer (49). Mutations in the *chvGI* region would also have to survive natural selection, outcompete other beneficial mutations that arose over the course of the experiment (clonal interference), and tolerate royal jelly (48). Therefore, observing mutations in the *chvGI* region across ten independent populations is remarkable and indicates the importance of this regulatory system and its impact on the bacterium’s ability to survive in royal jelly.

### ChvG and ChvI Regulatory Proteins

The ChvGI system was originally discovered in *A. tumefaciens* and *S. meliloti* (50–52). Although this system also functions in several free-living bacteria, it is particularly relevant in symbionts and pathogens, enabling acclimation to the stress manifested during intimate host association. ChvG is often regulated by the periplasmic protein designated ExoR, which contains multiple SEL1 (or tetratricopeptide, TPR) repeats (53, 54) known to mediate protein-protein interactions. ExoR interacts with the periplasmic domain of ChvG, negatively regulating its autophosphorylation activity, and thereby preventing phosphotransfer to ChvG. Stress conditions, commonly low pH, lead to proteolytic cleavage of ExoR and release of ChvG inhibition, conferring environmental control of the ChvG-ChvI regulon (55, 56).

These three proteins (ChvG, ChvI, and ExoR) are conserved regardless of whether the bacterium is known to interact with a host. Several Alphaproteobacteria have ChvG and ChvI orthologs, but do not have a protein similar to ExoR. This seems to be the case in *B. apis*, where there is a Sel1-repeat containing protein, but not one that is predicted to be secreted.

### Regulation of symbiosis in *Alphaproteobacteria*

By comparing *B. apis chvG* mutant behaviors to better understood ChvGI regulons, namely in *A. tumefaciens, S. meliloti* and *Rhizobium leguminosarum*, we can hypothesize how ChvGI regulates specific bacterial responses and the advantages this may confer in honey bee association. In other *Alphaproteobacteria*, ChvGI aids in the bacterial response to stress, likely envelope perturbation, and for symbionts, the transition from a free-living organism to a host-associated lifestyle (32). As observed in other *Alphaproteobacteria,* a mutant *chvG* or *chvI* can decrease host association (51, 57), a result that is consistent with our *chvG* mutants and their decreased ability to associate with honey bee larvae. We also observe diminished biofilm formation in a *chvG* mutant, which may relate to the mechanism behind the reduction we observed in larval association. ChvGI are also reported to regulate the production of exopolysaccharides, such as the complex branched polysaccharide succinoglycan. In *S. meliloti,* succinoglycan provides protection from the antimicrobial properties of plant-derived nodule-specific cysteine rich peptides (NCR), and mutations resulting in a hyperactive ChvGI system led to increased resistance to specific NCRs (58). In *B. apis* it is currently unclear if ChvGI also regulates exopolysaccharide production and the role this might play in aiding host association. Further study is required to fully understand the other requisite genetic mechanisms for *B. apis* to transition to a host-associated lifestyle.

Interestingly, in other systems, a mutated *chvG* leads to a decreased stress response and compromised overall cell viability when persisting in nutrient limitation or acidic environments (59). Royal jelly is a glandular secretion, produced by worker bees, that is extremely viscous, acidic, and contains antimicrobial peptides (60, 61). Curiously, in *B. apis*, it seems that a null *chvG* would be advantageous in an acidic environment such as royal jelly, suggesting that this system may function counter to other ChvGI systems.

We speculate that *B. apis* ChvGI may respond to a different pH since the ChvGI system in the zoonotic pathogen *Bartonella henselae* becomes activated when exposed to the physiological pH of blood (62). Beyond the null mutations in *chvG* that were extensively studied here, we also note frequent acquisition of point mutations within the *chvI* response regulator, particularly substituting from an arginine to a histidine (**Figure 3A**). Experimental evolution of *Escherichia coli* in fluctuating nutritional conditions and the ensuing pH changes consistently resulted in similar mutation from basic to acidic residues in a conserved transcription factor (63). It is tempting to speculate that these mutations may be significant for *B. apis* and its adaptation to pH fluctuations. Motility may be among the important responses to acid stress response in *B. apis.* As seen in other bacteria and symbioses, pH may act as a trigger for motility to promote *B. apis* movement towards an optimal pH, potentially sites it may colonize including the larval gut (64, 65). Since ChvGI is also known to regulate motility, it is possible that a null ChvG was selected because of its enhanced motility phenotype. However, at this juncture we cannot say for certain whether motility is a phenotype for which the royal jelly experimental evolution selected or if it is an indirect, collateral effect of a regulatory mutation.

### The link between ChvGI and flagellar regulation in *B. apis*

The link between motility and symbiosis has been well studied, particularly in its importance for the initial steps of host colonization where we see chemotaxis also plays a large role (36). ChvGI is known to regulate and impact motility, where an active ChvGI contributes to downregulation of genes related to motility and chemotaxis (66). We observe that a null *chvG* results in increased motility, with a higher abundance of flagellar-proteins and chemotaxis proteins overall (**Table S2**), consistent with how ChvGI regulate motility in other well-studied organisms (e.g. *A. tumefaciens* and *S. meliloti*). Strikingly, there is also a visible difference in the flagella produced by a *chvG* mutant relative to *B. apis* with a functional *chvG*. This led us to consider the basis for enhanced motility in JMM2 if both samples contain flagellar filaments. Our mass spectrometry results suggest that the ChvG null mutant JMM2 is producing a larger number of flagellar proteins overall, consistent with an increase in swimming behavior across the population of cells (**MOVIES S1, S2**). One possibility is that the JMM2 *chvG* mutant is producing a flagellar filament that A29 does not, as *B. apis* encodes two flagellin homologs, and we identified both distinct flagellins in the mass spectrometry analysis of sheared flagellar preparations for JMM2. It is also plausible that the regulation of flagellar filament production is impacted in the *chvG* null mutants, increasing the production of flagellar proteins overall and contributing to the subsequent increase in swimming behavior. However, we still question the thick protruding structures in the ChvG mutant, JMM2. Based on the high abundance of FlgK, a flagellar hook associated protein, these structures may be related to polyhook type appendages. Further studies will focus on the types and roles of these flagellar filaments to identify the role ChvGI plays in flagellation and motility in *B. apis* and how this may impact *B. apis* survivability in different hive micro environments.

### ChvGI broadly impacts the *B. apis* transcriptome

As in other bacteria, ChvGI has a clear impact on the transcriptome (34, 67, 68). Overall, none of the genes observed to change expression dramatically in the *chvG* null mutant stand out as clearly related to honeybee association nor the phenotypes that are impacted in the chvG mutant. However, a null *chvG* also led to a significant reduction in the gene *ctrA*. CtrA is a master cell cycle regulator known to play an important role in motility, chemotaxis, and cell division overall; most thoroughly studied in the alphaproteobacterium *Caulobacter crescentus,* CtrA is known to regulate the change between two distinct life cycles—swimming daughter cells and stalked, non-motile mother cells (41–43). CtrA is also thought to be important in the *S. meliloti* cell cycle as it differentiates into bacteroids during association with legume hosts (69). Although we observed a significant downregulation of *ctrA* in the transcriptome, there were no obvious cell cycle defects, growth changes, or morphological defects in our *chvG* null mutants. Either the downregulation observed is not significant enough to impact these aspects of microbial physiology or, as seen in several other *Alphaproteobacteria*, CtrA may act primarily as a regulator of motility in *B. apis* (70). This prompts questions regarding the lifestyle of *B. apis,* and how it may potentially change when host-associated. Further studies will focus on CtrA and its potential role in the symbiosis of *B. apis* and honeybees. Furthermore, the transcriptomic differences do not fully explain the drastic rescue of motility seen in our *chvG* null mutants. We hypothesize there may be other target genes, unidentified here, that may posttranscriptionally impact *B. apis* phenotypes.

### Conclusion

Our work underscores the significance of ChvGI as an important TCS in *Alphaproteobacteria* that regulates bacterial behavior in response to different types of stress and emphasizes the complexity of the molecular mechanisms that bacteria use when undergoing the many transitions involved in symbiosis. We specifically characterize ChvGI in a honey bee symbiont and identify the role it plays in mediating motility, biofilm formation, host association, as well as the fitness disadvantage it poses in our royal jelly selection. As a result, many questions remain regarding the importance of ChvGI in other microenvironments in the honey bee hive. Future experiments will aim to determine the ChvGI regulon and its potential to act as a contingency locus. This includes interrogation of the *B. apis* behavioral response in other stressful hive microenvironments it occupies to further our understanding of honey bee bacterial symbionts and the underlying genetic mechanisms they employ to withstand and transit through diverse honey bee environments.

## Data Availability

All sequences were deposited to NCBI including the complete *B. apis* A29 genome (PRJNA1427168) and the resequencing of evolved population (PRJNA1532902).

## Materials and Methods

### Growth and experimental evolution of Bombella apis A29

*B. apis* strain A29 was inoculated on Bacto-Schmehl (BS) (20) agar and grown overnight at 34°C, after which a single colony was used to inoculate 5 mL liquid cultures of BS broth at the same temperature, with shaking at 250 rpm. After an additional overnight incubation, 50 ul was transferred into each of the following treatments for a final dilution of 1:100: 10, 5-ml tubes of BS media (BS treatment) and 10, 5-ml tubes of a 50/50 mixture of BS broth and Stakich brand royal jelly (Larval diet/LD treatment). All resultant cultures were homogenized using a vortex mixer and incubated at 34°C without shaking. After another 24 hours, all cultures were thoroughly homogenized using a vortex mixer and 50 μl from each was transferred into BS broth and incubated at 34°C with 250 rpm shaking to permit recovery of CFUs. This regime was repeated for 28 days, with the BS treatment passaged in BS only with shaking at every other transfer, and the LD treatment passaged alternately between BS containing 50% royal jelly, and BS alone (with shaking). During passaging, one BS-treatment culture was contaminated, resulting in a final count of 19 experimentally evolved strains. At the end of 28 days, the resultant populations were used for population-level sequencing (see below) and subjected to single colony isolation to identify individual, isolated mutants. Individual mutations within the ChvG locus were confirmed for each isolate from the experimental population using PCR and Sanger sequencing. For all strains subjected to the royal jelly challenge, a culture at an OD_600_ of 0.4 was back diluted 1:10 into 3 mL of 30% royal jelly in BS broth. Every 6 hours, each culture was thoroughly homogenized, serially diluted in sterile PBS, and plated in triplicate on BS agar to count CFUs.

### Resequencing evolved populations

At the end of the experiment (day 28) entire populations for all replicates, and the ancestral A29 strain were grown up in BS broth and incubated at 34°C with 250 rpm shaking overnight. Overnight cultures were spun down to pellet cells, and DNA was extracted from cell pellets using the Qiagen DNeasy Blood & Tissue Kit. To reconfirm strain identity, and lack of contamination, we performed 16S PCR amplification and Sanger sequencing (via Quintara Biosciences).

Total DNA was submitted to SeqCoast Genomics for Illumina sequencing. Samples were prepared for whole genome sequencing using an Illumina DNA Prep tagmentation kit and unique dual indexes. Sequencing was performed on the Illumina NextSeq2000 platform using a 300-cycle flow cell kit to produce 2×150bp paired reads. 1-2% PhiX control was spiked into the run to support optimal base calling. Read demultiplexing, read trimming, and run analytics were performed using DRAGEN v3.10.11, an on-board analysis software on the NextSeq2000. Raw reads were downloaded from SeqCoast and QCs with Trimmomatic to remove adapters.

The Breseq pipeline was used to align reads to the B. apis A29 reference (PRJNA301156), call single nucleotide polymorphisms (SNPs) between strains, and identify putative new junctions (71). Because much of the B. apis genome is not functionally annotated, we used protein homology to annotate ChvGI using UniProt (72).

### Culturing bacteria

All *Bombella apis* strains were grown at 34°C, ambient oxygen, shaking at 230 rpm, in Bacto-Schmehl (BS) liquid media (20). BS liquid media is composed of 5% w/v D-glucose, 5% w/v D-fructose, 1% yeast extract, 4% v/v 5X Sigma M9 salt solution (M9956), and 0.2% v/v cation solution (1% [v/v]; 100 mM MgSO_4_ and 10 mM CaCl_2_ in diH_2_O), and 84.8% Milli-Q H_2_O. The cation solution must be autoclaved separately from the M9 solution. The final pH of BS media is 6.5.

### Biofilm assay

*B. apis* evolved strains and the progenitor, A29 were subjected to a crystal violet biofilm assay. For each isolate, a culture with an OD_600_ of 0.4 was used to make a 1:10 dilution in BS broth, across 8 technical replicates within a 96-well plate of BS broth, plus media-only controls. Cultures were allowed to grow overnight without shaking after which OD_600_ of each culture was measured before plates were washed by submerging in DI water twice and inverted on paper towels to dry. Once dry, 250 μl of crystal violet was added to each well and incubated at room temperature for 15 minutes. After incubation, the plate was again washed by submerging in water three times and inverted on paper towels to dry. Once dry, 250 μl of acetic acid was added to each well and incubated at room temperature for 15 minutes to dissolve the stained biofilm into a homogeneous solution. Finally, 200 μl of the acetic acid/crystal violet mixture was transferred to a new 96-well plate to measure absorbance at 550 nm (A_550_). Differences between the groups were assessed using a one-way ANOVA. Post-hoc pairwise comparison was performed using Tukey’s HSD in R 4.4

### Motility assay

To assess swimming motility, evolved *B. apis* strains and progenitor strain A29 were tested in BS swim plates containing 0.2% Difco agar. Petri plates were filled with 25 mL BS swim agar and solidified on the benchtop for 1 hour and 15 minutes. Once solidified, plates were inoculated from bacterial patches by using a pipette tip and stabbing halfway into the center of the agar and incubated at 34°C for 24 hours.

### Royal jelly challenge

Progenitor strain A29 and the 3 ChvG mutants were normalized to an OD_600_ of 0.4 and back diluted 1:10 into freshly prepared, UV sterilized, 30% royal jelly (Stakich brand royal jelly diluted with BS media) and left to grow for 24 hours at 34°C and shaking at 230 rpm. Growth was assessed by plating CFUs every 6 hours onto BS plates. Growth rate for each respective strain was calculated using the following formula: Ln(N_6_)-Ln(N_0_)/6, with N being the number of CFUs at time point 0 or 6.

### Larval collection and rearing

Larvae were collected from hives at Indiana University Research and Teaching Preserve Bayles Road field site in September 2024. To minimize the impact of genetic diversity, larvae were collected from the same colony.

Grafting and rearing protocols were executed according to Schmehl et al. 2011 and Parish et al. 2022 with the noted deviations. First and second-instar larvae were grafted from frames brought back to lab and stored in a 30°C room with 40% relative humidity (RH) using plastic grafting tools and placed into plastic queen cups in 48-well plates. To prevent contamination between larvae, queen cups were placed in every other well.

Larvae in the 48-well plates were then placed in plastic Tupperware with a damp paper towel to maintain humidity and incubated in darkness at 34°C and 75% RH.

All larvae were fed according to the diet recipes laid out in Schmehl et al., 2011. Larval diet was made fresh before each feeding and UV sterilized for 20 minutes to remove contaminants prior to feeding. LD was distributed midday daily to each larva in a flow hood using sterile pipettes.

The larval feeding schedule and bacterial inoculation occurred as follows:

On day zero freshly grafted larvae were fed 10 ul of larval diet A. 24 hours after grafting, larvae were fed 10 ul of larval diet A supplemented with 50 ug/mL of ampicillin to clear out bacteria from the hive environment. 48 hours post-grafting, larvae were fed 20 ul of larval diet B in addition to 10^5^ CFUs/mL of either A29 or JMM2 at mid-exponential phase, or the equivalent volume of sterile PBS. Larvae were only inoculated with bacteria once throughout the course of the experiment. The remaining days larvae were fed as follows: Larvae were fed 30 ul of diet C at the 72-hour mark, 40 ul of diet C 96 hours post-grafting, and 50 ul of diet C 120 hours after grafting. Larval diets are composed of varied amounts of royal jelly, glucose, fructose, and yeast extract as denoted in Schmehl et al., 2011. The brand of royal jelly used was: Stakich 100% Pure, Fresh All-Natural Royal Jelly.

On day 6, 144 hours after the initial grafting, living larvae were washed in the following steps: embryo wash, 10% bleach, embryo wash, and PBS. Embryo wash contains 70 g NaCl w/v and 5 mL of Triton v/v for 250 mL.

Differences between the groups were assessed using a Kruskal-Wallis test. Post-hoc pairwise comparison was performed using a Wilcoxon rank sum test with a Benjamini-Hochberg false discovery rate correction method in R 4.4.

### RNAseq

WT and JMM2 were grown in BS media overnight and back diluted 1/10 the following day. RNA was harvested once both cultures reached mid-exponential phase (OD_600_= ∼0.6) using the PureLink RNA Mini kit following the manufacturers’ instructions. rRNA was removed using Terminator reaction buffer A. Quality of the RNA was then assessed on the TapeStation and sent to SeqCoast genomics for 150 bp paired end Illumina sequencing. Briefly, reads were aligned using Bowtie2, counted using HTseq-count, assessed for differential expression using Deseq2, and COGclassifier to determine COG groups. The volcano plot and the bar graph showing COG classification were generated using R 4.4.

### Mass spectrometry of flagellar filaments

Preparation of flagellar filaments for mass spectrometry was done as previously described for *A. tumefaciens* (38). Progenitor strain A29 and one ChvG mutant representative, JMM2, were grown overnight at 34°C, subcultured into 500 mL of BS media until they reached mid exponential phase. Cultures were then centrifuged at 10,000 x g for 15 minutes to remove cells. Strain JMM2 was centrifuged for an additional 10 minutes as this strain is difficult to pellet. The supernatant for each respective sample was collected and ultracentrifuged at 74,500 x g for 30 minutes at 4°C in a fixed-angle Beckman 70 Ti rotor. The pellet was resuspended in 20 mL of phosphate-buffered saline (PBS), and centrifuged at 17,500 x g for 10 minutes to remove any remaining cells. The resulting supernatant was finally ultracentrifuged once more at 74,500 x g for 30 minutes to concentrate the flagellar preparation. The remaining pellet was resuspended in 100 ul of PBS and sent for LC-tandem mass spectrometry. Samples were resuspended and denatured in 8 M urea with 100 mM ammonium bicarbonate, pH 7.8. Disulfide bonds were reduced by incubation for 45 min at 57°C with a final concentration of 10 mM Tris (2-carboxyethyl) phosphine hydrochloride (Sigma Aldrich). A final concentration of 20 mM iodoacetamide (Sigma Aldrich) was then added to alkylate these side chains and the reaction was allowed to proceed for one hour in the dark at 21 °C. Samples were diluted to 1 M urea using 100 mM ammonium bicarbonate, pH 7.8. A total of 0.5 µg trypsin (Promega) was added, and the samples were digested for 14 hours at 37 °C. Individual samples were desalted using ZipTip pipette tips (EMD Millipore), dried down and resuspended in 0.1% formic acid. Samples were analyzed by LC-MS on an Orbitrap Fusion Lumos equipped with an Easy Nano-LC1200 HPLC (ThermoFisher Scientific). Buffer A was 0.1% formic acid in water. Buffer B was 0.1% formic acid in 80% acetonitrile. Peptides were separated on a 30-minute gradient from 0% B to 3% B. Precursor ions were measured in the Orbitrap with a resolution of 60,000. Fragment ions were measured in the Orbitrap with a resolution of 7,500. The spray voltage was set at 1.8 kV. Orbitrap MS1 spectra (AGC 1×10^6) were acquired from 350-2000 m/z followed by data-dependent HCD MS/MS (collision energy 30%, isolation window of 2 Da) for a three second cycle time. Charge state screening was enabled to reject unassigned and singly charged ions. A dynamic exclusion time of 30 seconds was used to discriminate against previously selected ions. The LC-MS/MS data was searched against the Uniprot *B. apis* strain A29 proteome (downloaded 05 2026) using Proteome Discoverer (v2.5.0.400) note: this is the information for that additional analysis we performed with the updated proteome.

Trypsin was set as the protease specificity allowing for two missed cleavages. Carbamidomethylation of cysteine residues was set as a static modification. Oxidation of methionine, pyroglutamine formation on peptide-terminal glutamine, acetylation of the protein amino terminus and loss of the protein amino terminal methionine residue was set as dynamic modifications. The precursor mass tolerance was 10 ppm and the fragment ion tolerance was 0.04 Da. Error rates were determined using Percolator with an FDR rate of 0.01.

### Motility movies

Strains were grown to mid-exponential phase in BS media and diluted into minimal BS for optimal imaging conditions. Phase contrast movies were acquired on a Nikon Eclipse Ti2 microscope with a 100x objective and processed using NIS-Elements software (Nikon).

### Electron Microscopy

Four samples of *B. apis*: A29-Swim, JMM2-Swim, A29-Broth, and JMM2-Broth, were prepared for TEM by negative staining. This was done by placing 10 µl of the sample, diluted 1/10 in dH_2_O, on glow discharged (Pelco easiGlow, Ted Pella, Inc., Redding CA), carbon coated 300 mesh copper grids (Catalog number CF300-Cu, Electron Microscopy Sciences, Hatfield, PA) for 5 minutes. The sample droplet was removed by wicking with Whatman No. 1 filter paper and replaced with 10 µl of a 2% aqueous solution of uranyl acetate (Catalog number 22400, Electron Microscopy Sciences, Hatfield, PA). The stain was removed with filter paper, and the grids dried for five minutes. The grids were viewed and photographed with an HT of 80 KV with a JEOL company, JEM 1010 transmission electron microscope equipped with a Gatan company, Multiscan camera, model 794. The image size was 1024 x 1024. For SEM, four 12 mm diameter coverslips (Catalog number 26020, Ted Pella, Inc., Redding, CA) were coated with 0.1% poly L lysine (Catalog number 19320-B, Electron Microscopy Sciences, Hatfield, PA). This was done by placing 10 µl drops of PLL on each of the coverslips for 5 minutes. Excess PLL was then removed by a dH_2_O rinse. 10 µl of each sample of *B. apis*: A29-Swim, JMM2-Swim, A29-Broth, and JMM2-Broth, were placed on their own prepared coverslip for 5 minutes. Excess sample was removed by drawing the drop away with filter paper, and the coverslip was placed in 2.5% glutaraldehyde (Catalog number 16020, Electron Microscopy Sciences, Hatfield, PA) in 0.1 M sodium cacodylate buffer, pH 7.2 (Catalog number 12300, Electron Microscopy Sciences, Hatfield, PA). Coverslips were rinsed in dH_2_O, dehydrated in a graded ethanol dehydration series, critical point dried (Samdri-790, Tousimis Research Corporation, Rockville MD), and sputter coated with a 12 nm layer using a 80% gold/ 20% palladium target (CCU-010 sputter coater, Safematic GmbH, Zizers, Switzerland). Coverslips were affixed to aluminum SEM stubs (Catalog number 16145, Ted Pella, Inc., Redding, CA) using carbon double stick tape (Catalog number 77816, Electron Microscopy Sciences, Hatfield, PA). Samples were imaged with a Teneo VS SEM (Thermo Fisher Scientific Inc., Waltham, MA). Images were 4096 x 3536 pixels, taken at an HT of 5 kV, specimen current of 50 pA, WD of 8.3 mm, and 1 µs dwell time.

